# Rhizoliths identified as prehistoric filing tools for fishhook production on San Nicolas Island, California

**DOI:** 10.1101/2021.01.05.425479

**Authors:** Sebastian K.T.S. Wärmländer, Kevin N. Smith, René L. Vellanoweth, Ryan Moritz, Kjell Jansson, Tim Gooding, William E. Kendig, Sabrina B. Sholts

**Affiliations:** Division of Biophysics, Stockholm University, Sweden; Cotsen Institute of Archaeology, UCLA/Getty Conservation Programme, UCLA, USA; Department of Archaeology and Classical Studies, Stockholm University, Sweden; Dept. of Anthropology, UC Davis, USA; Dept. of Anthropology, California State University Los Angeles, USA; Dept. of Materials and Environmental Chemistry, Stockholm University, Sweden; Dept. of Mineral Sciences, National Museum of Natural History, Smithsonian Institution, Washington, DC, USA; Dept. of Anthropology, National Museum of Natural History, Smithsonian Institution, Washington, DC, USA

**Keywords:** Electron microscopy, X-ray diffraction, Residue analysis, Surface chemistry, Ancient technology, California archaeology, Native American, California Channel Islands, Nicoleño culture

## Abstract

Chemical analysis of archeological objects can provide important clues about their purpose and function. In this study, we used scanning electron microscopy (SEM) and chemical spectroscopy (SEM-EDS and XRD) to identify a white residue present on cylindrical rhizoliths from a component at an archaeological site (CA-SNI-25) on San Nicolas Island, California, dated ca. AD 1300 to 1700. The residue was found to consist of biogenic calcite and aragonite particles, different in composition and morphology from the CaCO3 particles in the rhizoliths, but identical to marine shell material. These results, together with observations of surface micro-wear patterning on fishhooks and rhizoliths, replicative experiments, *in* situ spatial analysis, and other archaeological evidence, show that rhizoliths were used as files in a larger tool kit for crafting shell fishhooks. Our findings shed new light on the technological innovations devised by Native Americans to exploit the rich marine resources surrounding the Channel Islands, and provide the first analytical evidence for the use of rhizoliths as a production tool.

## 1. Introduction

Even with modern technology it can be challenging to elucidate the original functions of archaeological objects. Yet, information regarding the manufacture, use, trade, and discard of items encountered in the archaeological record is key to understanding past societies. In North America, few places preserve more details of human prehistory than the California Channel Islands (Figs. 1 and S1–S2), where the inhabitants have left a near continuous record of occupation for the last 13,000 years (Rick et al., 2005; Erlandson, J. M. et al., 2008; Erlandson et al., 2011; Byrd and Raab, 2007). On these islands, archaeologists have documented evidence of a thriving maritime culture that extracted food and raw materials from island and offshore resources: stone and bone tools were used to craft a range of objects such as reed and wood-plank canoes, twined and coiled basketry, and shell beads and other ornaments (Erlandson et al., 2011; Braje et al., 2005; Wärmländer et al., 2011; Gamble, 2002; Meighan and Eberhart, 1953; Gifford, 1947; Bleitz, 1993). What remains less clear, however, are the functional relationships between items found in Channel Island sites. How, for instance, were the expedient flakes, abraders, burins, and other artifacts used in the past? What were their specific functions and how were they incorporated into tool and ornament production sequences?

**Fig. 1.**
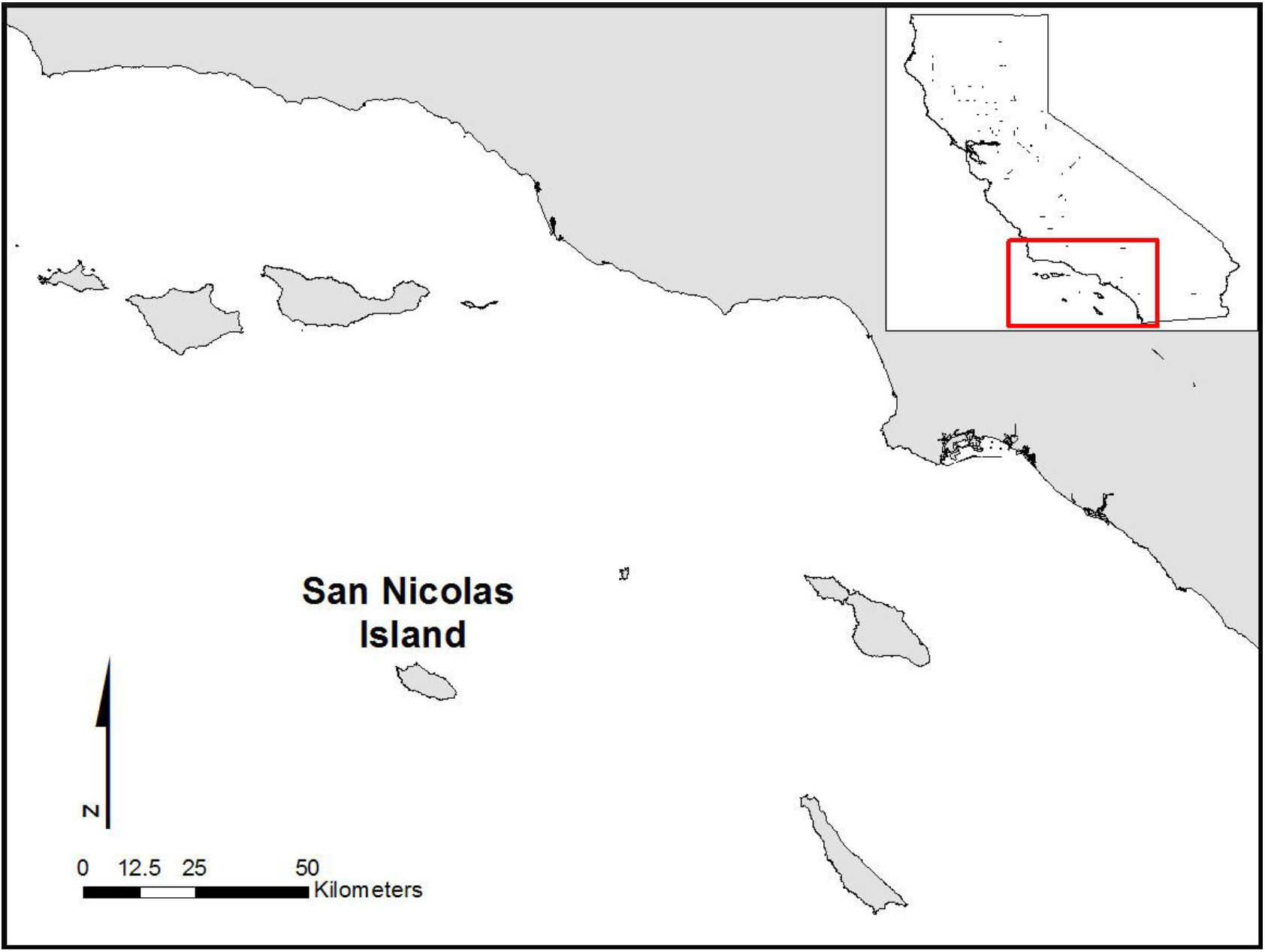
Map showing the location of San Nicolas Island.

In this study, we investigated one such class of artifact, namely the relatively ubiquitous rhizoliths that are found in numerous Channel Island sites. Rhizoliths, or rhizoconcretions, are natural concrete root casts, often cylindrical in shape, with a hollow core and typically ranging from ~3-8 cm long when found in archaeological contexts. These concretions form when dissolved calcium carbonate precipitates around plant roots and solidifies as gritty cylinders with bumps, nodes, and nodules reminiscent of the roots that formed them (Fig. 2) (Stewart and Thorson, 1994). On the Channel Islands rhizoliths are often found eroding out of raised beaches (Fig. S3). Although they typically display a rough surface with inclusions of quartz grains (i.e., sand), rhizoliths excavated from archaeological sites on San Nicolas Island often display smooth surfaces with occasional patches of an unknown white residue (Fig. 2). Almost all rhizoliths with such modifications have been found at human occupation sites, suggesting that their distinctive features are related to human activities. This is somewhat puzzling, as rhizoliths have not been previously shown to be involved in any human technology, although some researchers have proposed that rhizoliths could have been used as crafting tools (Bleitz and Salls, 1993; Kendig et al., 2010; Steele, 2006; Smith, 2013).

**Fig. 2.**
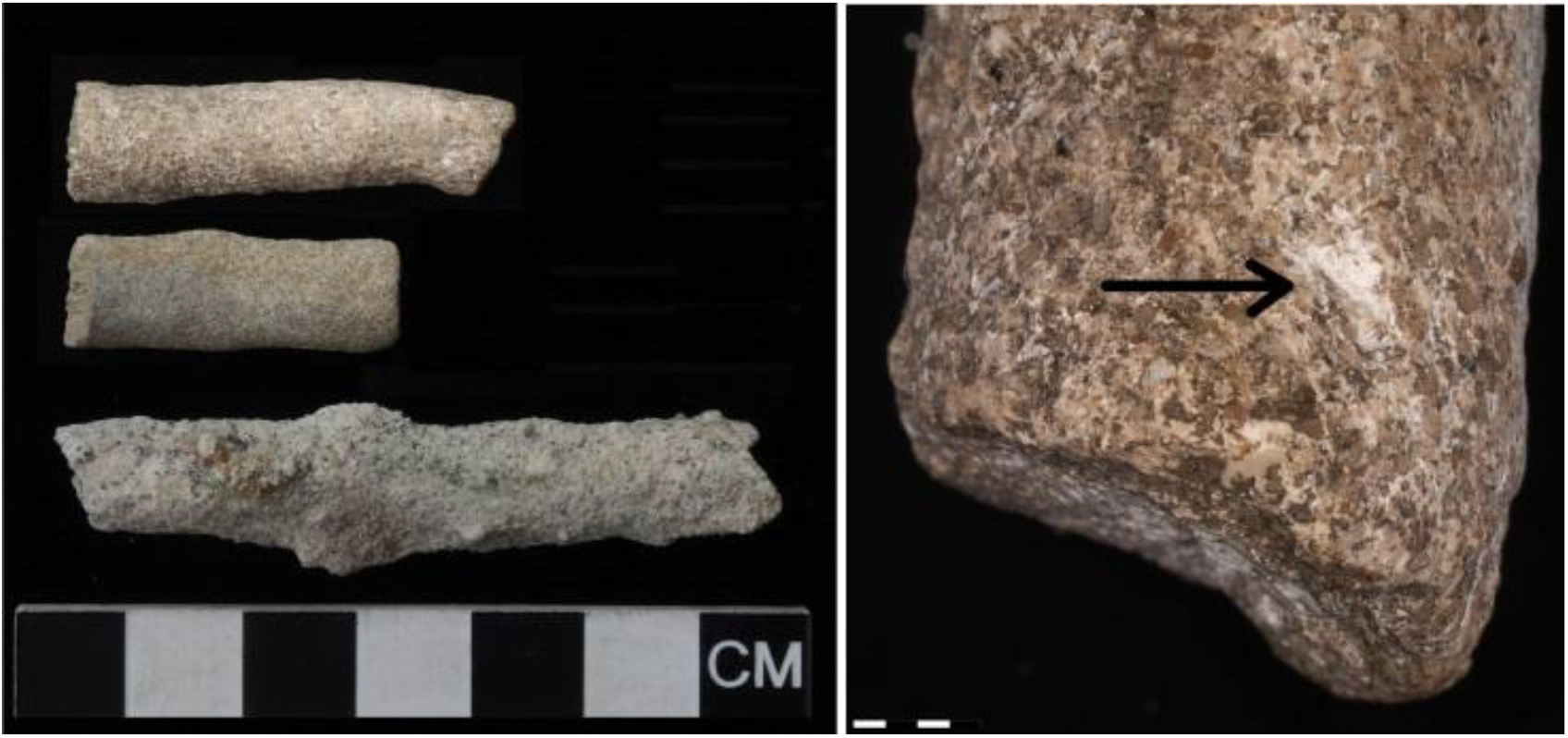
Left: Two rhizoliths with smoothed surfaces excavated at the Tule Creek site on San Nicolas Island (specimen 5038, top; specimen 5039, middle) and one modern rhizolith with a natural rough surface (bottom). Right: Close-up of the white surface residue on archaeological rhizolith 5038 (indicated by arrow).

Using chemical spectroscopy and electron microscopy, we analyzed residue patches and other surface features of two rhizoliths excavated from the Tule Creek site on San Nicolas Island. Combining the analytical data with information from the archaeological record, replicative experiments, and additional analyses of modern comparative materials, we were able to understand how native San Nicolas Islanders incorporated rhizoliths as production tools in their fishing technology.

## 2. The Tule Creek site (CA-SNI-25), San Nicolas Island

Almost 100 km from mainland California, San Nicolas Island is the most remote of the eight California Channel Islands (Figs. 1 and S1) (Kendig et al., 2010). This semi-arid island covers an area of about 80 km^2^, and is sparse in terrestrial resources. It contains only a few perennial fresh water springs, low growing shrubs, and terrestrial animals that include the white-footed deer mouse *(Peromyscus maniculatus),* side-blotch lizard *(Uta stansburiana),* island night lizard *(Xantusia riversiana),* southern alligator lizard *(Elgaria multicarinata),* island fox *(Urocyon littoralis),* and land snail *(Micrarionta feralis)* (Schoenherr et al., 1999). On the other hand, San Nicolas Island is surrounded by the richest kelp beds in the Channel Islands region, and this unique ecosystem harbors an abundance of marine life (Browne, 1994; Engle, 1994). The island’s prehistoric inhabitants, the Nicoleño, caught fish and sea mammals using a variety of strategies, involving single and composite bone gorges and hooks, circular and J-shaped shell fishhooks, harpoons, and nets (Bleitz, 1993; Mariani, 2001; Meighan, 1954; McKenzie, 2007). ^14^C-dates of archaeological deposits as well as finds of lithic crescents, a flaked stone technology dating to the Terminal Pleistocene/Early Holocene, indicate an early occupation of the island at least 8,000 years ago (Davis et al., 2010). Archaeological evidence of large village sites, communal cemeteries, and increased inter-island and cross-channel trade (Martz, 2005) show that the populations of San Nicolas and the other Channel Islands grew significantly in size and social complexity during the Late Holocene (roughly 3,500 years ago until European contact) (Arnold, 2001; Gamble, 2008; Kennett, 2005).

The Tule Creek site (CA-SNI-25) is a relatively intact village with minimal subsurface disturbance (Fig. S2), located 3.2 km southeast of Thousand Springs, the northernmost point on San Nicolas Island. The site overlooks Corral Harbor, one of the few safe canoe anchorages on the island. Over 75 radiocarbon dates indicate three discrete occupational time periods for this site: 4800-4300 cal BP, 2800-1350 cal BP, and 650-250 cal BP. CA-SNI-25 exhibits archaeological evidence of domestic as well as ceremonial activities, including a broad array of lithic reduction concentrations, fish and shellfish processing areas, dog burials (Bartelle et al., 2010; Vellanoweth et al., 2008), artifact caches, two balancing stone features (Knierim et al., 2013), hearth alignments, and pit features that may be associated with feasting events. The site’s abundant artifacts and exposed features caught the attention of antiquarians and relic hunters early in the post-contact history of the island. In the early twentieth century, Rogers (Rogers, 1930) described the site as containing numerous house pits, eleven communal houses (nine of which had been used as cemeteries), several sandstone saws and fishhooks, and a vast quantity of flaked stone.

Beginning in 2001, recent excavations have concentrated primarily on two loci, designated as East Locus and Mound B. Hundreds of human-made objects from the latest occupation period (AD 1300 - 1700) have been unearthed, including ornamental shell artifacts, fishhooks in various stages of production, and stone assemblages including arrow points, drills, knife blades and spear points. Some items were produced from non-local materials, such as steatite from Santa Catalina Island, obsidian from Coso and Obsidian Butte, chert from Monterey and San Miguel Island, as well as chalcedony, clear rock crystal, and fused shale from unidentified mainland sources (Erlandson et al., 1997; Erlandson, Jon M. et al., 2008; Rick et al., 2001; Arnold, 1987; Rosenthal, 1996; Cannon, 2006). This demonstrates the existence of a well-established prehistoric inter-island and cross-channel trade network (Arnold, 1990). The vast majority of artifacts identified were however produced from locally available materials such as stone, bone, and shell (Meighan and Eberhart, 1953; Cannon, 2006; Guttenberg, 2014; Gifford, 1947). Geologically, San Nicolas Island consists primarily of uplifted and alternating Eocene sandstones, siltstones, and conglomerates (Schoenherr et al., 1999; Meighan and Eberhart, 1953; Vedder and Norris, 1963). This composition provides a highly indurated and extremely hard sandstone (Kendig et al., 2010) that was used by the Nicoleño to make bowls, pestles, and the type of artifact that Rogers in 1930 described as a “stone saw” (Rogers, 1930). These “saws” appear to have been used as abrading tools primarily in the manufacture of the island’s characteristic circular shell fishhooks (Kendig et al., 2010). At Tule Creek, the sandstone abraders have often been found together with rhizoliths, suggesting a possible connection between the two types of objects (Kendig et al., 2010).

## 3. Materials and Methods

Ongoing field work at CA-SNI-25 has so far yielded a total of 203 rhizolith pieces from East Locus. Of these pieces, 143 display smoothed surfaces indicative of use-wear, and six display traces of white surface residue. Two such rhizoliths, with inventory numbers 5038 and 5039, were selected for chemical analysis (Fig. 2). In addition, a handful of modern rhizoliths were collected from the island for comparative purposes. The excavations on San Nicolas Island, which is US federal land, were carried out under a permit issued by the US Navy to author RLV in accordance with the US Archaeological Resources Protection Act (ARPA). The samples were analyzed at the National Museum of Natural History (Smithsonian Institution), Washington, D.C., and at Stockholm University, Sweden, in compliance with all relevant legislation and ethical guidelines. Rhizoliths 5038 and 5039 are currently curated at the Department of Anthropology, California State University in Los Angeles (CSULA), together with other material excavated from San Nicolas Island.

Scanning electron microscopy (SEM) images and energy-dispersive spectrometry (EDS) data were recorded for small scrapings of residue particles, shell/rhizolith material, and complete fishhooks, using various SEM units: a Nova NanoSEM 600 (FEI, USA) and a JSM-7000F (JEOL Ltd., Japan), both equipped with field emission guns, and a table-top Hitachi TM-3000 (Hitachi Ltd., Japan). All samples were analyzed without carbon coating at low kV and low beam current to avoid charge build-up. A 0.5 x 0.5 mm cross-section of modern red abalone shell was prepared using argon beam polishing under a zero-degree angle to the surface, employing a JEOL SM-09010 Cross Section Polisher (JEOL Ltd., Japan) running at 5 kV for 12 hrs. X-ray microdiffractograms of minute sample particles were recorded on a D/max-Rapid micro-XRD unit (Rigaku Corp., Japan) operating at 50kV/40mA with an Mo anode, and analyzed using JADE v.9 software (Materials Data Inc., USA). The spatial distributions of excavated rhizoliths and fishhooks at the Tule Creek site were analyzed using the ArcMap 10.2 software in the ArcGIS suite (ESRI, Redlands, CA, USA).

## 4. Results and discussion

The SEM-EDS analysis of the white residue on the archaeological rhizoliths (Figs. 3 and S4) identified Ca, C, and O in approximate ratios of 1:1:3, suggesting that the residue is calcium carbonate (CaCO3). This was confirmed by XRD analysis of small scrapings of the residue, which identified two CaCO3 phases: calcite and aragonite (Figs. 4 and S5). The presence of calcite in the residue might not be surprising, as the XRD and SEM-EDS analyses of modern rhizoliths identified calcite as their main component, with some inclusions of quartz (SiO2) particles (Figs. 3, 4, and S5). The general absence of aragonite in the rhizolith bulk material, however, indicates an exogenous source for the aragonite observed in the residue.

**Fig. 3.**
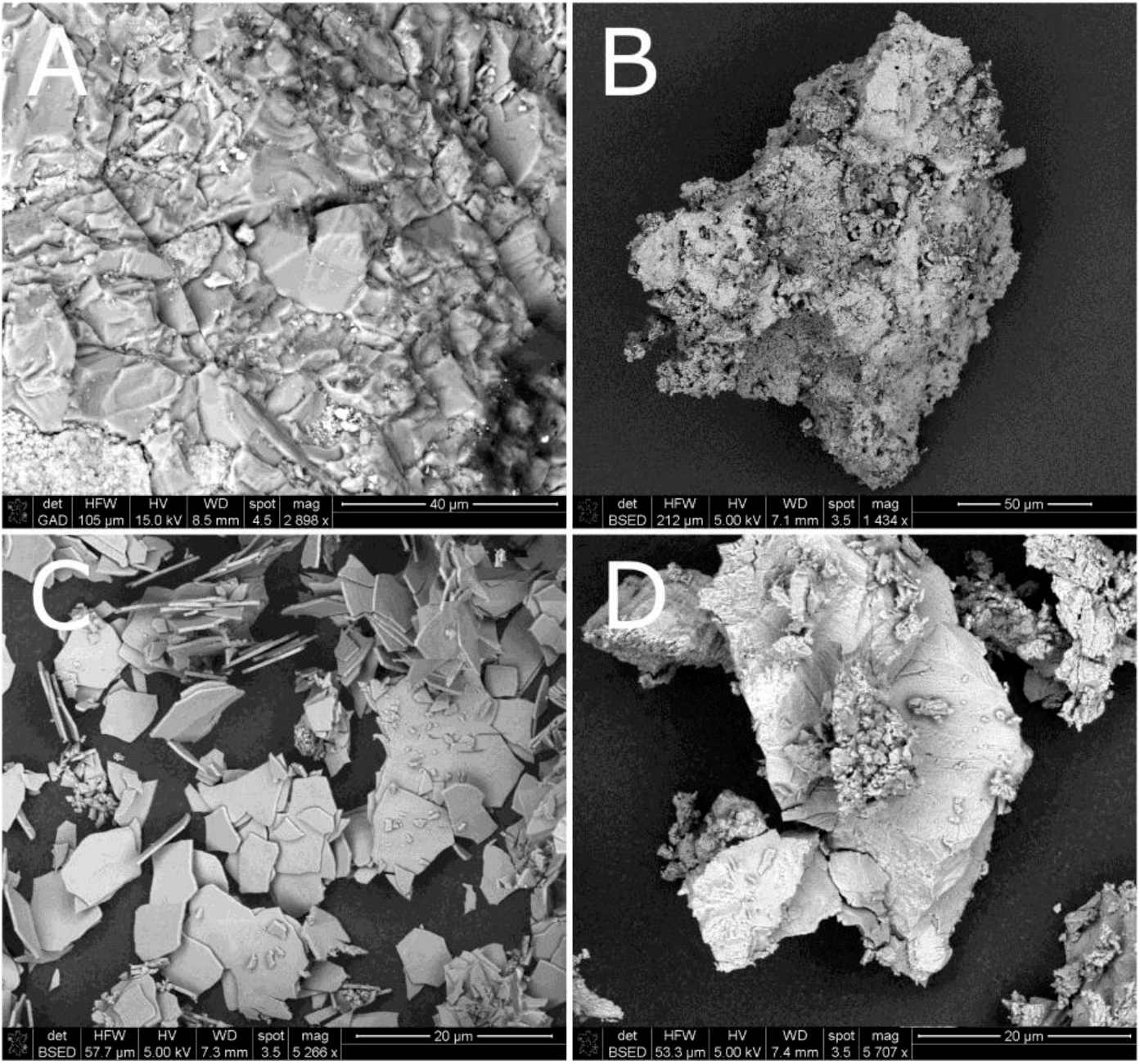
SEM backscatter images of A) white residue from archaeological rhizolith 5038; B) calcite particles from a modern rhizolith; C) aragonite plates from the inner nacreous layer of red abalone shell; D) calcite particles from the outer epidermis layer of red abalone shell.

**Fig. 4.**
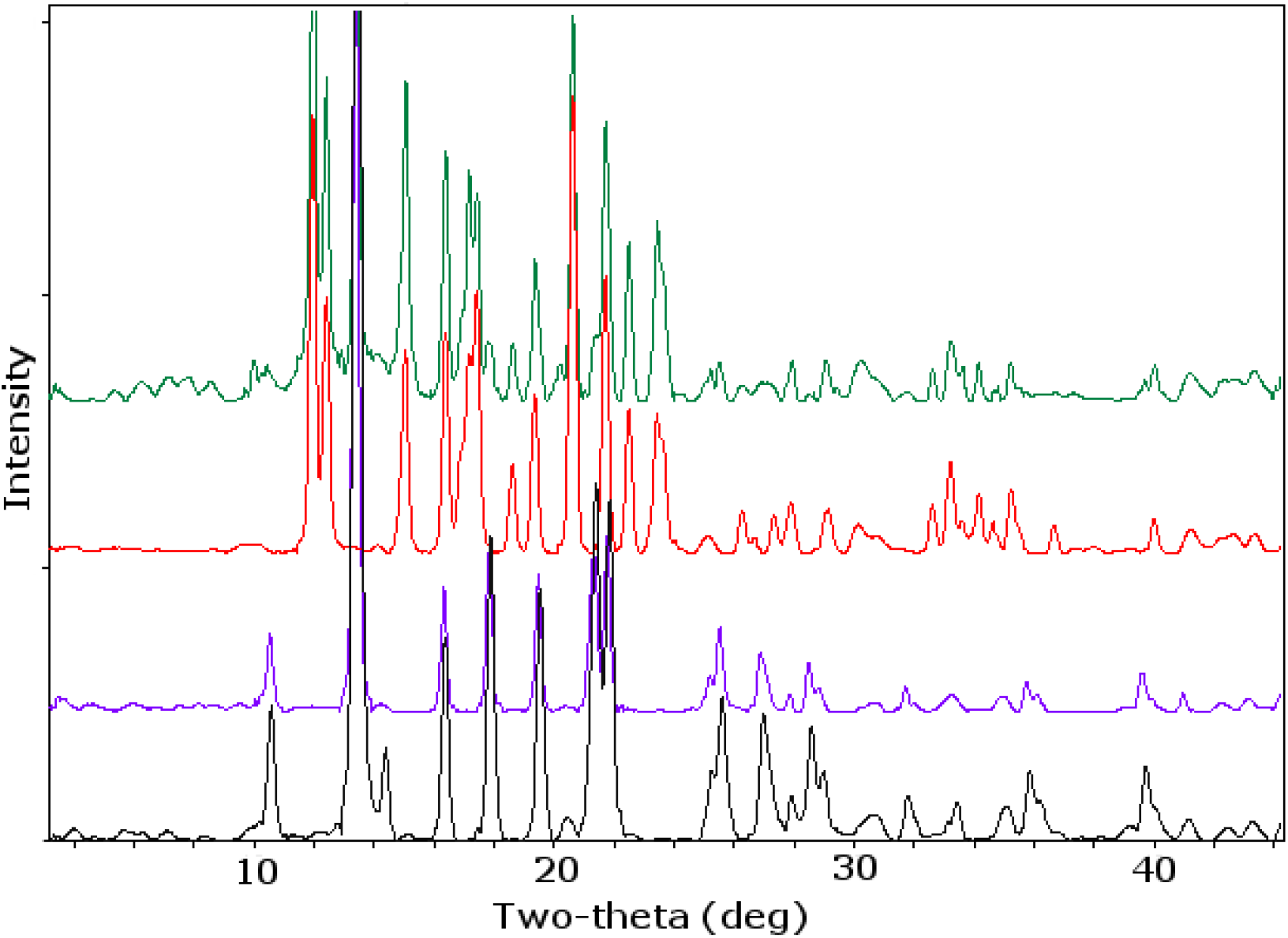
X-ray microdiffractograms of minute scrapings of particles from: a modern rhizolith (black, calcite); outer epidermis layer of abalone shell (purple, calcite); inner nacreous layer of abalone shell (red, aragonite); residue from archaeological rhizolith no 5038 (green, aragonite and calcite). Additional information is provided in Figs. S5 and S6.

Aragonite occurs naturally in certain geological formations in California (Coleman and Lee, 1962), but the most readily available source of this mineral is the nacreous (mother-of-pearl) layer in the shells of certain marine mollusks along the Pacific coast, such as red abalone (*Haliotis rufescens*; Fig. 5). The inner nacreous layer of abalone shell consists of characteristic aragonite plates, 10–20 μm wide and around 0.5 μm thick (Fig. 3C). The abalone’s outer epidermis layer consists of boulder-shaped calcite particles organized in layers that likely reflect biological growth patterns (Figs. 3D, S4, and S6). These calcite particles are morphologically identical to the residue particles from the archaeological rhizoliths (Figs. 3A, 3D, and S4), suggesting that the residue’s calcite particles also originate from marine shells and not from the rhizoliths themselves.

**Fig. 5.**
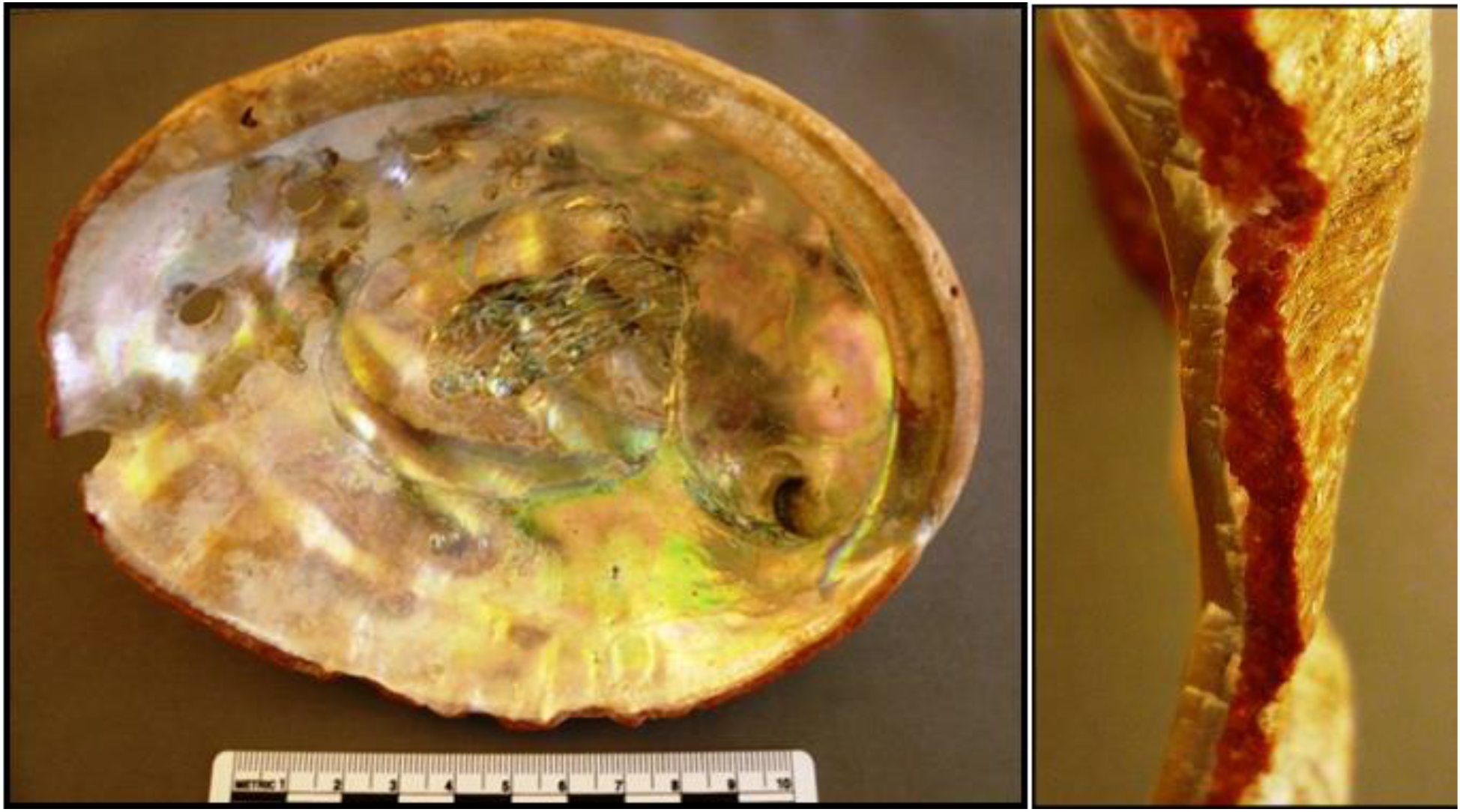
Photographs of a modern red abalone shell (*Haliotis rufescens*). Left: The interior nacre (mother-of-pearl). Right: Cross-section showing the inner nacreous layer and the outer red epidermis layer.

Although particles of sea shells are occasionally present in the eroded soil from which the rhizoliths form (see Fig. S7 for small fragments of aragonite plates in the matrix of a modern rhizolith), the proportion of aragonite in the residue is much higher. Thus, the current results unambiguously clarify the residue’s origin in marine shells, and also confirm the usefulness of SEM-EDS paired with micro-XRD analysis for virtually non-destructive identification of surface traces of unknown inorganic material (Scott et al., 2009; Smith et al., 2018; Roos et al., 2020; Smith et al., 2015; Bartelink et al., 2015), even in cases where the trace and the bulk material both consist of calcium carbonate.

In addition to residue patches, the smoothed areas of the archaeological rhizoliths also display patterns of surface microstriations, as is evident in SEM images (Fig. 6). Together with conchoidal fractures in quartz particle inclusions (Fig. 6), this clearly demonstrates that mechanical forces have been applied along the surface plane. Given the results of the chemical analysis, these forces likely relate to abrasive activities involving marine shells. The Channel Islanders harvested huge quantities of abalones and other shellfish from rocky intertidal and subtidal habitats in the waters along the island shorelines, and their shells were used extensively for a range of purposes (Vellanoweth and Erlandson, 1999; Erlandson, J. M. et al., 2008). At the Tule Creek site, the archaeological record contains a large number of circular fishhooks and fishhook preforms - in varying stages of manufacture - made from abalone shell. These fishhooks are often found together with worn rhizoliths (Kendig et al., 2010), as shown in the find distribution maps for the excavation units at Tule Creek (Fig. 7). The spatial correlation is established by high numbers of rhizoliths and fishhooks in the northern and central units, and low numbers especially in the south-west corner of the excavation area. Thus, the combined archaeological and chemical data suggest that rhizoliths might have been connected with production of fishhooks from marine shells. This idea has previously been suggested by some archaeologists (Kendig et al., 2010; Bleitz and Salls, 1993; Steele, 2006), but has so far not been properly tested.

**Fig. 6.**
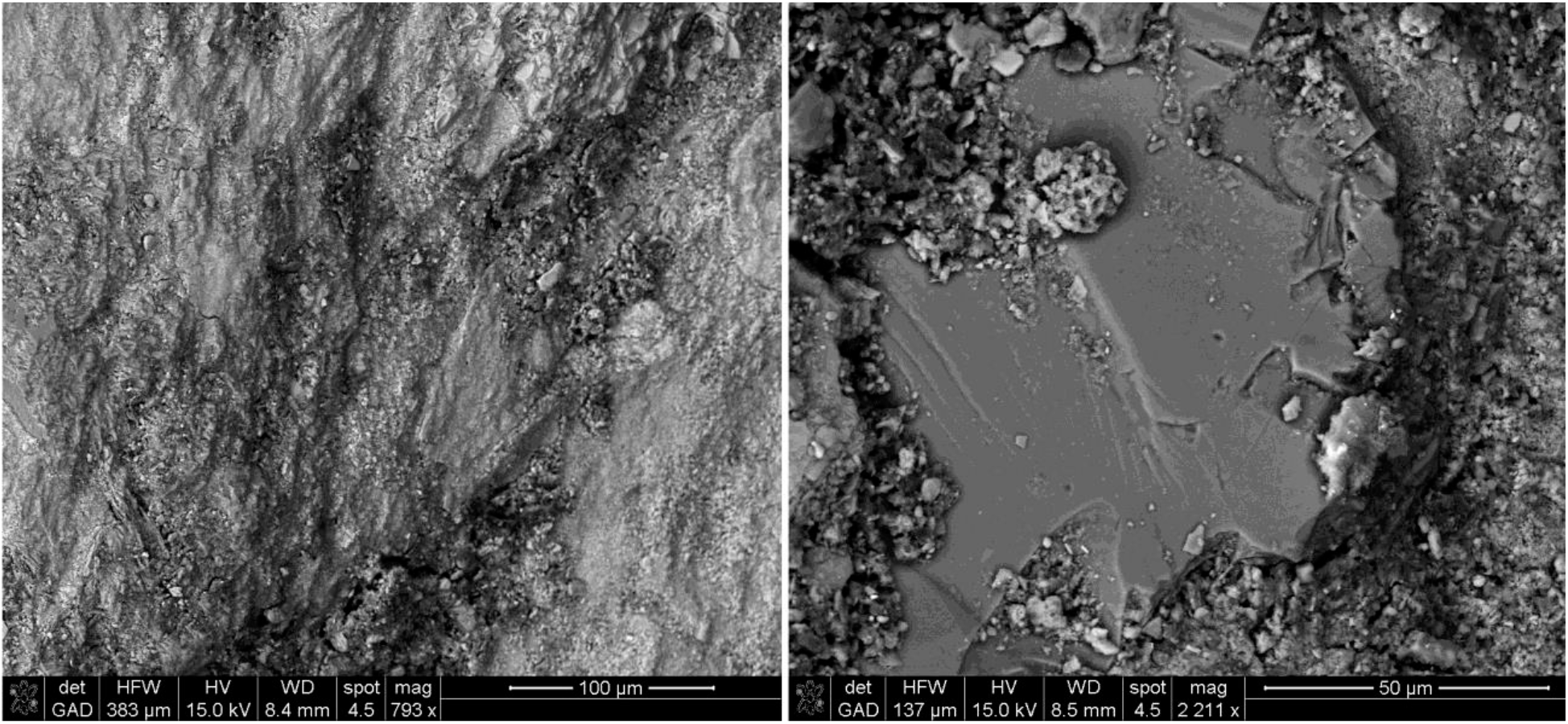
SEM backscatter images of archaeological rhizolith 5039, which has a smoothed surface. Left: surface topography showing microstriations. Right: quartz grain inclusion displaying a conchoidal fracture parallel to the surface plane, showing that mechanical forces have been applied in that direction.

**Fig. 7.**
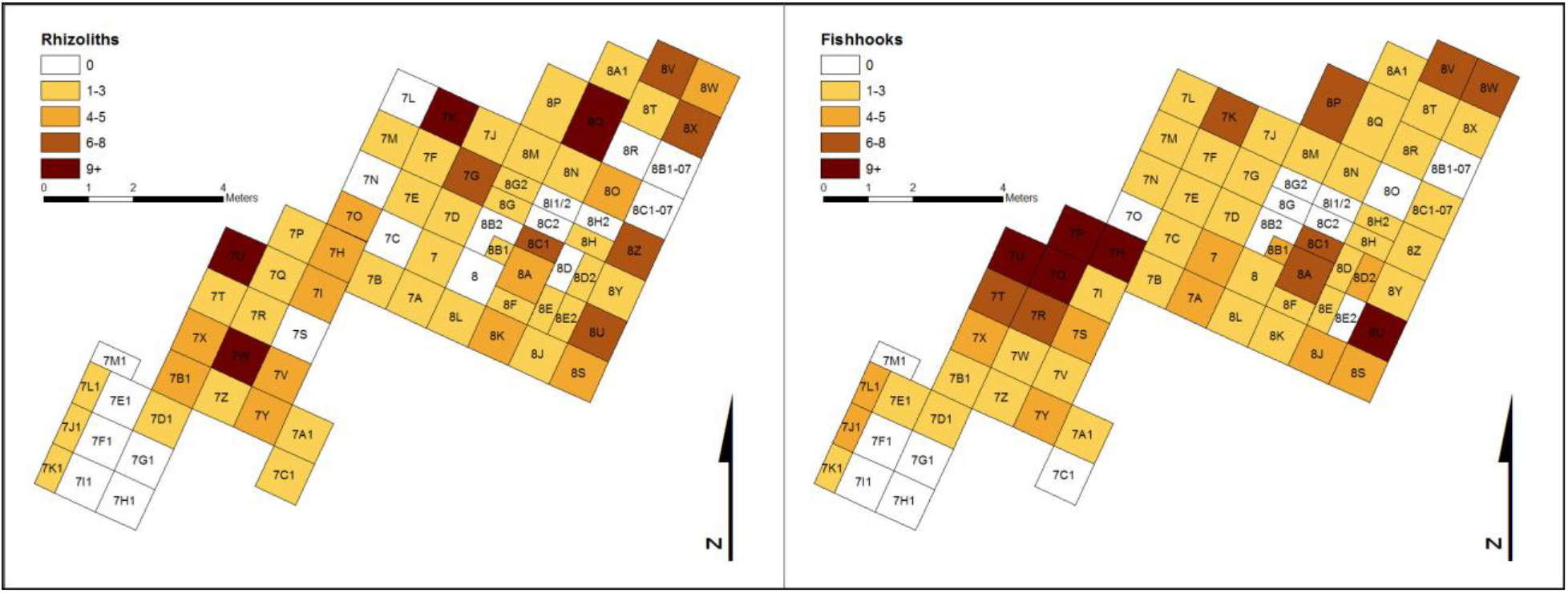
Maps showing the distribution of excavated rhizoliths (left) and fishhook fragments (right) at East Locus of the Tule Creek site on San Nicolas Island.

To investigate whether and how rhizoliths could have been employed in fishhook manufacture, we conducted replicative studies using modern rhizoliths from San Nicolas Island and abalone shell from the California coast (Smith, 2013; Smith et al., 2018; Smith et al., 2015). Red abalone was collected along the coastline of Sonoma and Mendocino counties, where stocks are still abundant and less threatened than in southern California, and crafted into fishhooks using traditional tools and techniques (Smith, 2013). Replicative studies of this type have been successfully employed previously to improve the understanding of prehistoric Channel Islands technology (Nigra and Arnold, 2013; McKenzie, 2007; Kendig et al., 2010). Our results show that rhizoliths can be effectively used as “rat-tail” files to shape, expand, and refine the inner curvature of the circular fishhooks common to San Nicolas Island (Figs. 8 and S8). During filing, the modern rhizoliths obtained smoothed surfaces similar to those present on the archaeological rhizoliths at the Tule Creek site (Figs. 2 and S9). The experimental rhizoliths regularly broke after one or two fishhooks had been refined, leaving two fragments behind: the distal worn portion and the proximal non-worn handle portion. Both types of fragments are common at the Tule Creek site, which further supports the hypothesis that the archaeological rhizoliths were used as filing tools in this manner. These findings also suggest that rhizolith files were “expedient tools,” i.e. implements typically produced with minimal time or effort from abundant and easily accessible local materials, and generally used and discarded without retooling (Kendig et al., 2010; Andrefsky Jr., 2005).

**Fig. 8.**
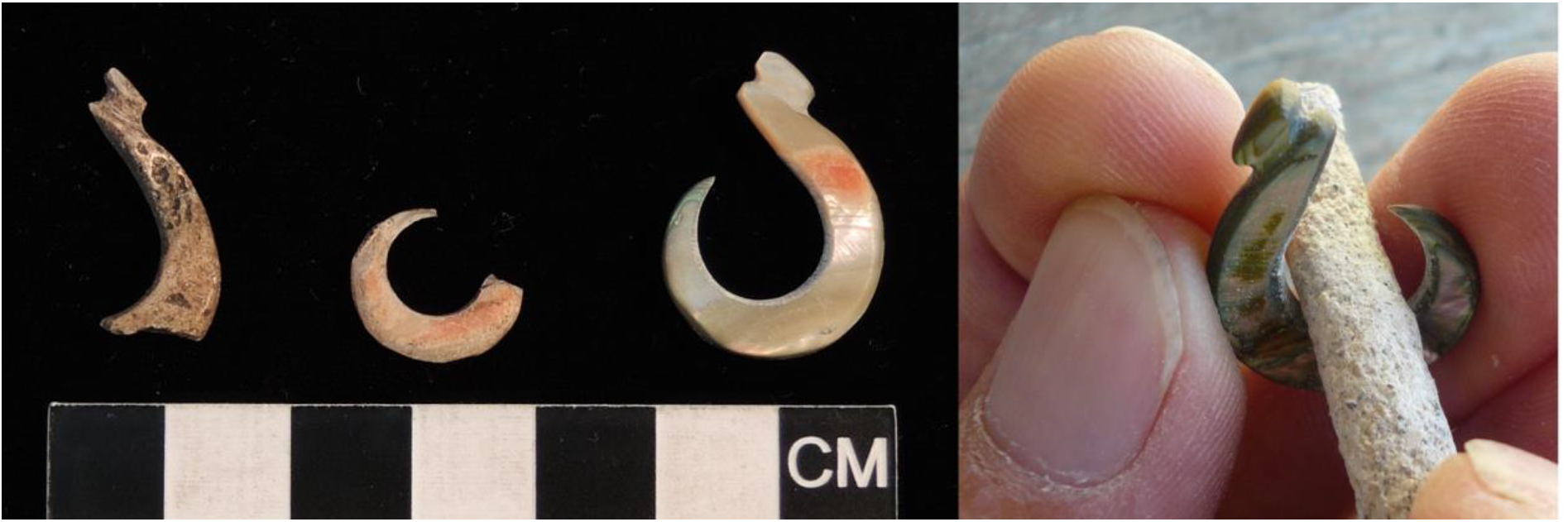
Left: Two archaeological fragments of circular fishhooks excavated at the Tule Creek site on San Nicolas Island (#4042, left; #711, center), together with a modern replica (right). All three hooks are made from nacre (mother-of-pearl). Right: The rhizolith functions as a “rat-tail” file for enlarging and refining the inner curvature of the hook. Note how the filing also wears down the rhizolith, giving it the same smoothed appearance as archaeological rhizoliths.

The replicative experiments furthermore allowed us to characterize the tool marks left by rhizoliths on shell fishhooks. Close-up SEM images of the inner curve of a modern replicated fishhook show an area covered with shallow markings originating from rhizolith filing (Figs. 9A and S10). Corresponding SEM images of an archaeological fishhook reveal several markings consistent with rhizolith filing on the side of the hook and, to a lesser extent, in the inner curve (Figs. 9B and S10). These observations clearly support the idea of rhizolith filing being a step in the manufacturing process. They also suggest that both the inner curve and the sides were filed, although most original tool marks have likely been lost due to wear and weathering from use and/or post-depositional processes. Moreover, the archaeological fishhooks are almost exclusively made from shell nacre, which has unusual material properties: the many layers of aragonite plates, held together by elastic biopolymers, form a lamellar structure that is strong, resilient, and inhibits transverse crack propagation. These properties are illustrated in SEM images of an abalone shell cross-section (Figs. 9C-D and S11–12). While the outer calcitic part displays multiple cracks running in different directions, the nacreous part shows cracks running into it and coming abruptly to a halt due to the material’s structure. Individual aragonite plates may be brittle, but broken plates in one layer do not reduce the strength of the plates in the next layer (Figs. 9C-D and S11–12). The layered microstructure, however, is not equally strong in all directions, which might explain why the sides of the fishhooks display better preservation of tool marks, as they are here imprinted parallel to the aragonite layers. The overall material toughness was likely the rationale for making fishhooks from nacre instead of the calcitic shell layer, which is harder but more likely to fracture and break. In the debate over fishhook material choices, Tartaglia (Tartaglia, 1976) has argued that it was the intriguing iridescent colors of nacre that were desired (just like for most other items made from this material), as nacre hooks might have worked similar to modern fishing lures. Yet, it is believed that the hooks were used with baits, and McKenzie (McKenzie, 2007) saw no beneficial effect of nacre iridescence when he compared the effectiveness of baited and unbaited shell fishhook replicas in fishing experiments.

**Fig. 9.**
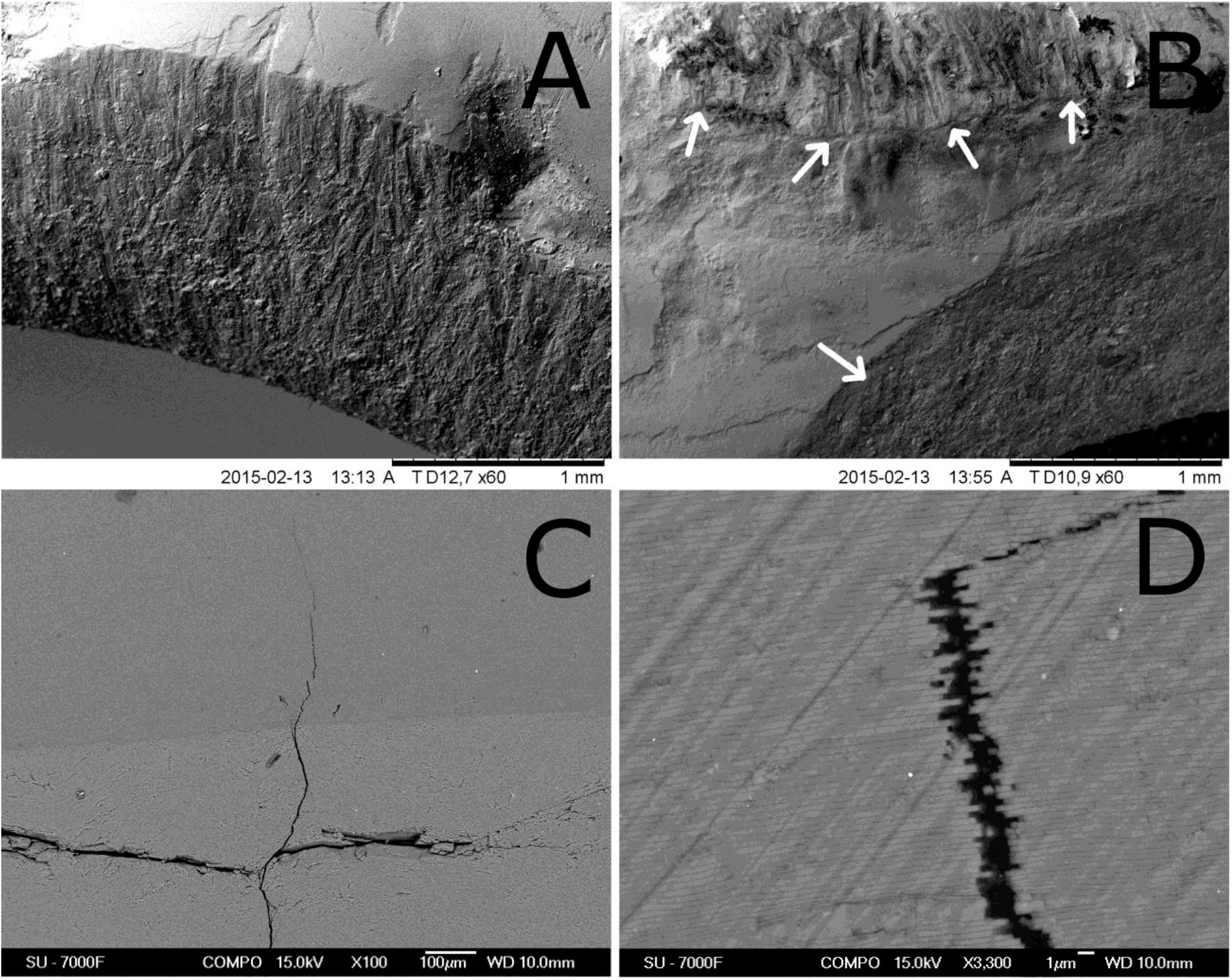
SEM backscatter images of shell fishhooks (top) and abalone shell material (bottom). A) The inner curve of a modern replicate fishhook. Multiple tool marks from filing with a rhizolith are present. B) Side and inner curve of archaeological fishhook #711. Multiple tool marks are observed on the side of the hook, and some also on the inner surface (indicated by arrows). C) Cross-section of a red abalone shell showing the intersection between the calcitic part (bottom) and the nacreous part (top) – compare Fig. 5. The calcitic part displays multiple cracks that stop upon entering the nacre. D) Close-up (x3300) of the crack in the nacreous layers of aragonite plates, showing that the crack consists of a series of missing or damaged aragonite plates.

As all available evidence suggests that rhizoliths were used as filing tools in shell fishhook production, the results of this study may help clarify archaeological finds at the Nursery Site (CA-SCLI-1215) on nearby San Clemente Island. Here, two rhizoliths (one of them previously misidentified as a degraded mammal bone and later re-analyzed by KNS as part of this study (Bleitz and Salls, 1993; Smith, 2013)) were found in a pouch of woven sea-grass primarily containing fishing implements ^14^C-dated to around AD 500 (Bleitz and Salls, 1993). If the two rhizoliths indeed were tools in this prehistoric “tackle box,” as seems likely, then together these two sites show that rhizoliths were employed in fishhook manufacture on the southern Channel Islands for a period of at least 1,000 years. The use of rhizoliths could, however, be as old as the circular single-piece shell fishhook itself, which first appeared on the Channel Islands around 3,000 years ago as an important complement to the earlier composite bone fishhooks and bone gorges (Rick et al., 2002). The single piece hook captures fish *via* mouth rather than gut hooking (Robinson, 1942), and its characteristic incurvature (Figs. 8 and S8) makes it less likely to snag when angling in kelp beds and rocky reefs (Robinson, 1942), which allowed the Channel Islanders to add larger quantities of fish from such habitats to their diet. In this view, the findings presented here expand our understanding of how the Nicoleño used local materials and technological innovation to exploit the rich marine resources around the California Channel Islands (Rick et al., 2005). Although many marine foods are lower ranked than terrestrial resources, their diversity and predictable abundance were arguably crucial for many early groups of modern humans (Marean et al., 2007; Parkington, 2001; Erlandson and Braje, 2011; O’Connor et al., 2011; Marean, 2014; Parkington, 2003; Klein and Steele, 2013; Steele, 2010). Recent studies indicate that maritime economies developed already 42,000 years ago in East Timor (O’Connor et al., 2011), and that oceanic navigation supported by marine foods likely facilitated the earliest peopling of the Americas (Erlandson and Braje, 2011; Erlandson et al., 2011; Erlandson et al., 2007). In terms of human adaptation to the California Channel Islands, the shell fishhook was a key technology for marine subsistence, on par with the water-bottle basket (Braje et al., 2005) and the plank canoe (Gamble, 2002).

In other prehistoric coastal locales, fishhooks appear to have been crafted using abrading/filing tools from various other materials, such as coral fragments on Polynesian Henderson Island (Weisler, 1994), large tropical sea urchin spines on Isla Espíritu Santo in the Gulf of California, Mexico (Fujita, 2012), shark skin on the northern Channel Island of Santa Cruz (Arnold and Graesch, 2001), and slate files at Tecolote Canyon on the mainland coast near Santa Barbara (Rick and Erlandson, 2008). Although these suggested material choices have so far not been confirmed with analytical chemistry techniques such as SEM-EDS or XRD, the two latter findings imply - if correct - a large regional and cultural variation in fishhook production tools already within the California Channel Islands region, possibly related to differences in local resources. From an economic and risk management perspective, it would be preferable to use ever-present local materials for basic subsistence technology, rather than traded goods with less reliable availability. Nonetheless, we have previously shown that innovations in lithic technology could diffuse rapidly across North America (Sholts et al., 2012), and under suitable conditions (i.e. plants growing in calcareous soil experiencing wet/dry cycles) rhizoliths form naturally in many places including the other Channel Islands (Stewart and Thorson, 1994). Even though it appears that the earliest Channel Islanders experienced little gene flow from the mainland (Sholts et al., 2010), neither San Nicolas nor any other Channel Island was ever culturally or economically isolated from the mainland or other islands during prehistoric times (Erlandson et al., 1997; Erlandson, Jon M. et al., 2008; Rick et al., 2001). Future research will hopefully clarify to what extent the use of rhizoliths as filing tools spread beyond San Nicolas Island.

The replicative experiments (Smith, 2013) together with previous research (Kendig et al., 2010; Smith et al., 2015; Smith et al., 2018) indicate that rhizolith filing was only one stage of a multistep manufacturing process for fishhook production. By the same token, rhizoliths were only one part of a multi-component fishhook manufacturing toolkit. Understanding the specific tools and steps involved in prehistoric fishhook production was one of the main questions raised by the first archaeologists and antiquarians to work in the Channel Islands region, and it has been a longstanding issue ever since (Schumacher, 1875). Ongoing research employing the type of chemical analysis presented here is expected to identify the complete set of production tools employed on San Nicolas Island, which arguably included coarse sandstone abraders and reamers (Kendig et al., 2010; Smith et al., 2015; Smith et al., 2018) and possibly also chert drills and meta-volcanic pecking tools (Smith, 2013).

## 5. Conclusions

In this work, we present results from chemical analyses, archaeological excavations, and replicative experiments that strongly support the idea that rhizoliths were used as files during the production of circular shell fishhooks on San Nicolas Island, California. While no single result is conclusive in itself, as a whole the evidence is compelling. We believe that combined use of these three approaches will be very useful in future archaeological research. To the best of our knowledge, this is the first study presenting analytical evidence for rhizoliths being used as manufacturing tools. This finding thereby expands the existing repertoire of natural resources employed in ancient lithic technologies.

## Acknowledgments

Funding and support for the archaeological fieldwork was provided by Humboldt State University, California State University, Los Angeles, and the U.S. Navy (Naval Base Ventura County and NAVAIR, Point Mugu). We thank Lars Eriksson at Stockholm University for help with the chemical analyses, Lauren Mirasol at CSULA for helping with logistics, and Wendy Teeter and Lucius Martin at the Fowler Museum at UCLA for their help in accessing and reanalyzing the Nursery Site fishing kit.

## Supplementary material

**Supplementary Figure S1.**
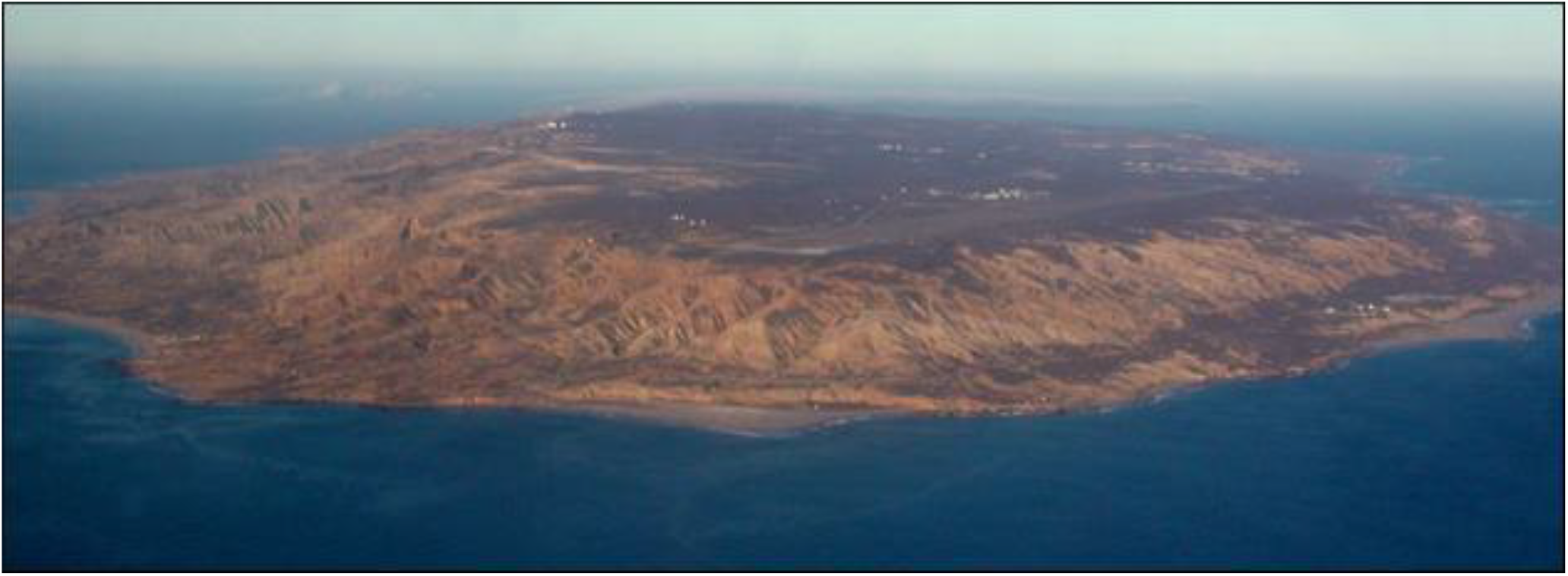
Aerial photograph of San Nicolas Island.

**Supplementary Figure S2.**
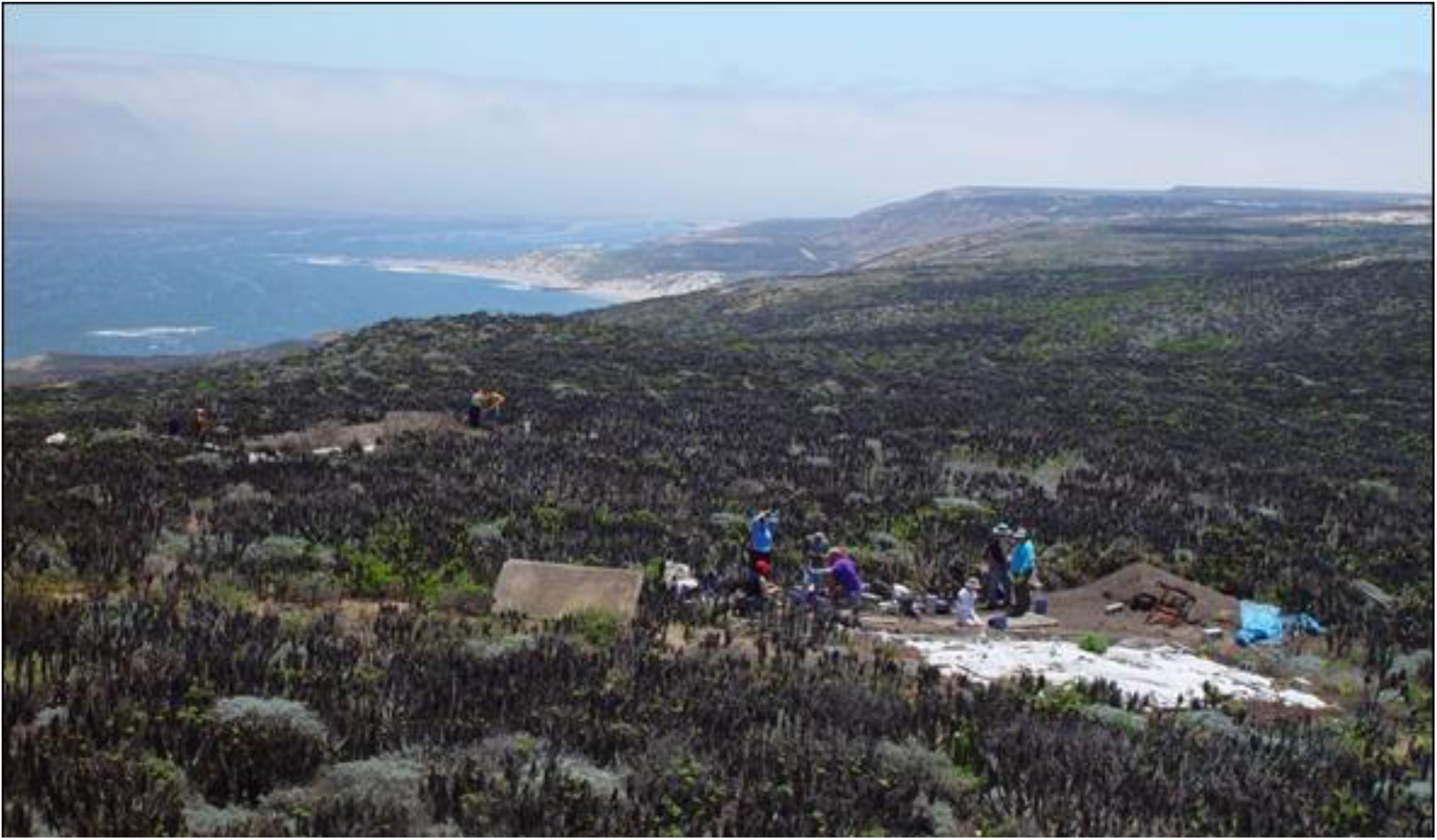
The Tule Creek site (CA-SNI-25) on San Nicolas Island, occupied ca. AD 1300 - 1700, showing evidence of ceremonial and domestic activities in the East Locus (left) and domestic activities in Mound B (right).

**Supplementary Figure S3.**
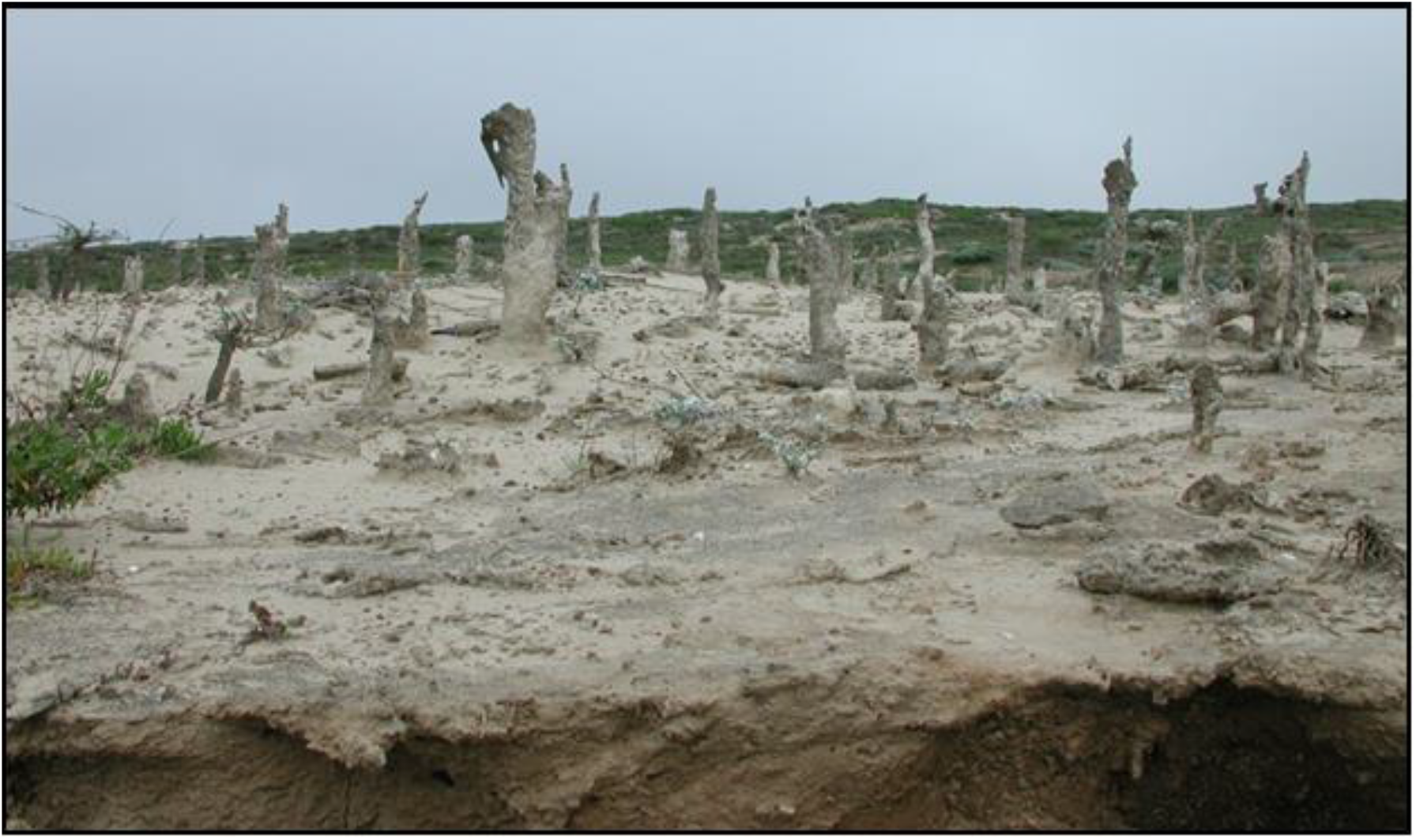
Rhizolith “forest” on San Nicolas Island. The concretions form when dissolved calcium carbonate precipitates and hardens into a tube around the roots of plants growing in a dune or shell midden. The “forests” are created by erosion forces, which remove the uppermost sand layer and thereby expose the rhizoliths.

**Supplementary Figure S4.**
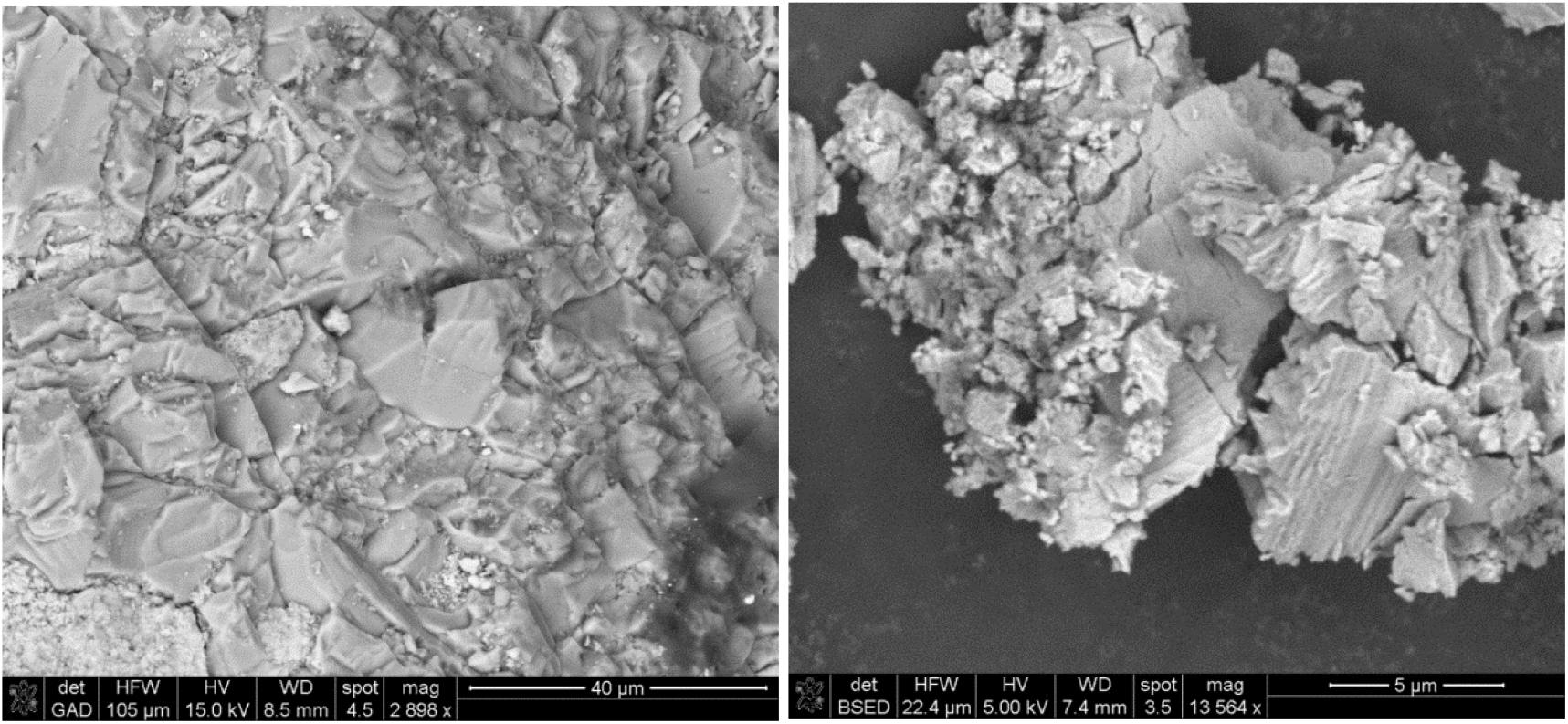
SEM images of CaCO3 residue from an archaeological rhizolith (left) and biogenic calcite particles from the outer layer of an abalone shell (right). Note the similar morphologies of the particles.

**Supplementary Figure S5.**
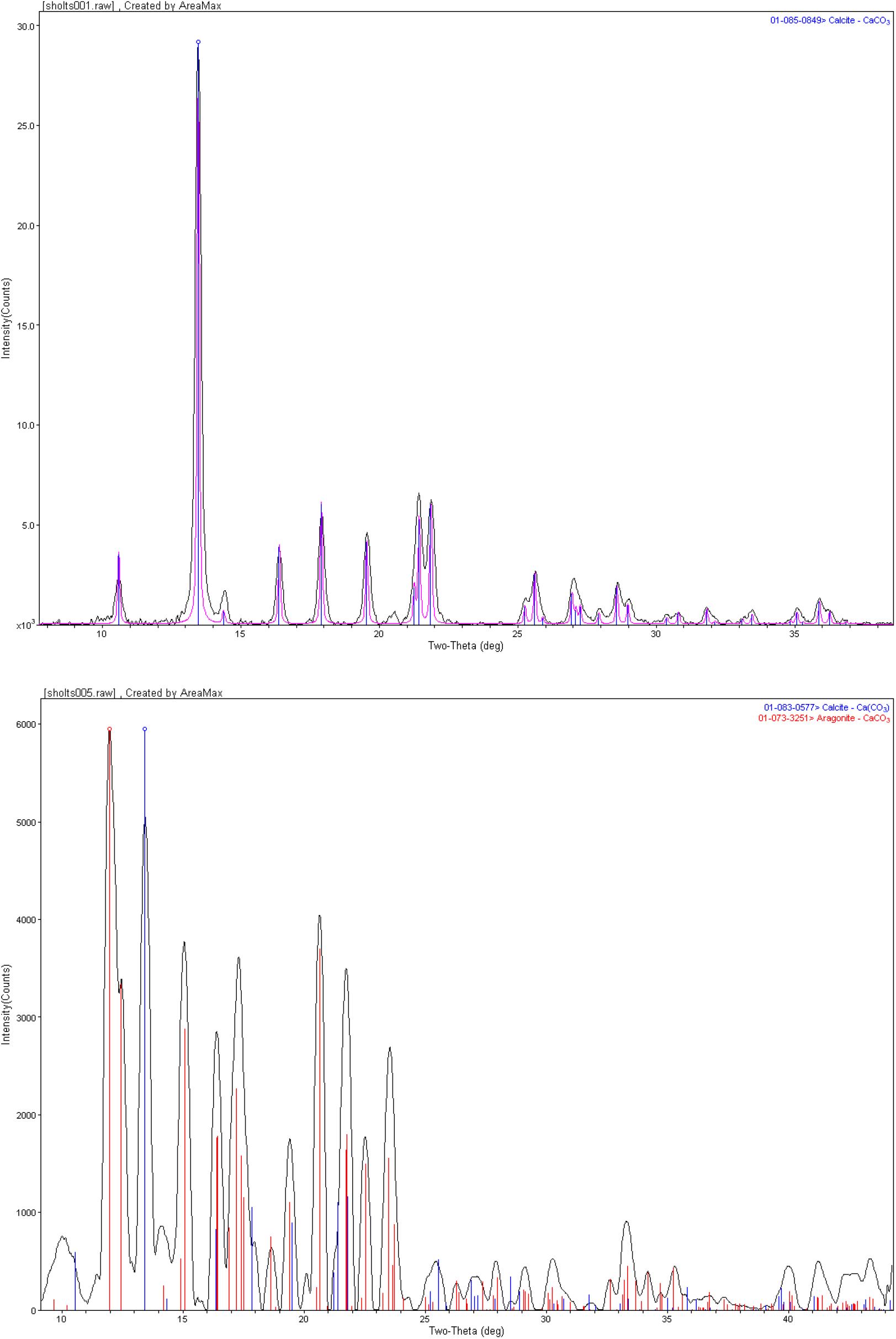
X-ray diffractograms of scrapings from a modern rhizolith (top) and white residue from archaeological rhizolith 5038 (bottom), perfectly matching reference data for respectively calcite (ICDD # 01-085-0849) and both calcite and aragonite (ICDD # 01-083-0577 and 01-073-3251).

**Supplementary Figure S6.**
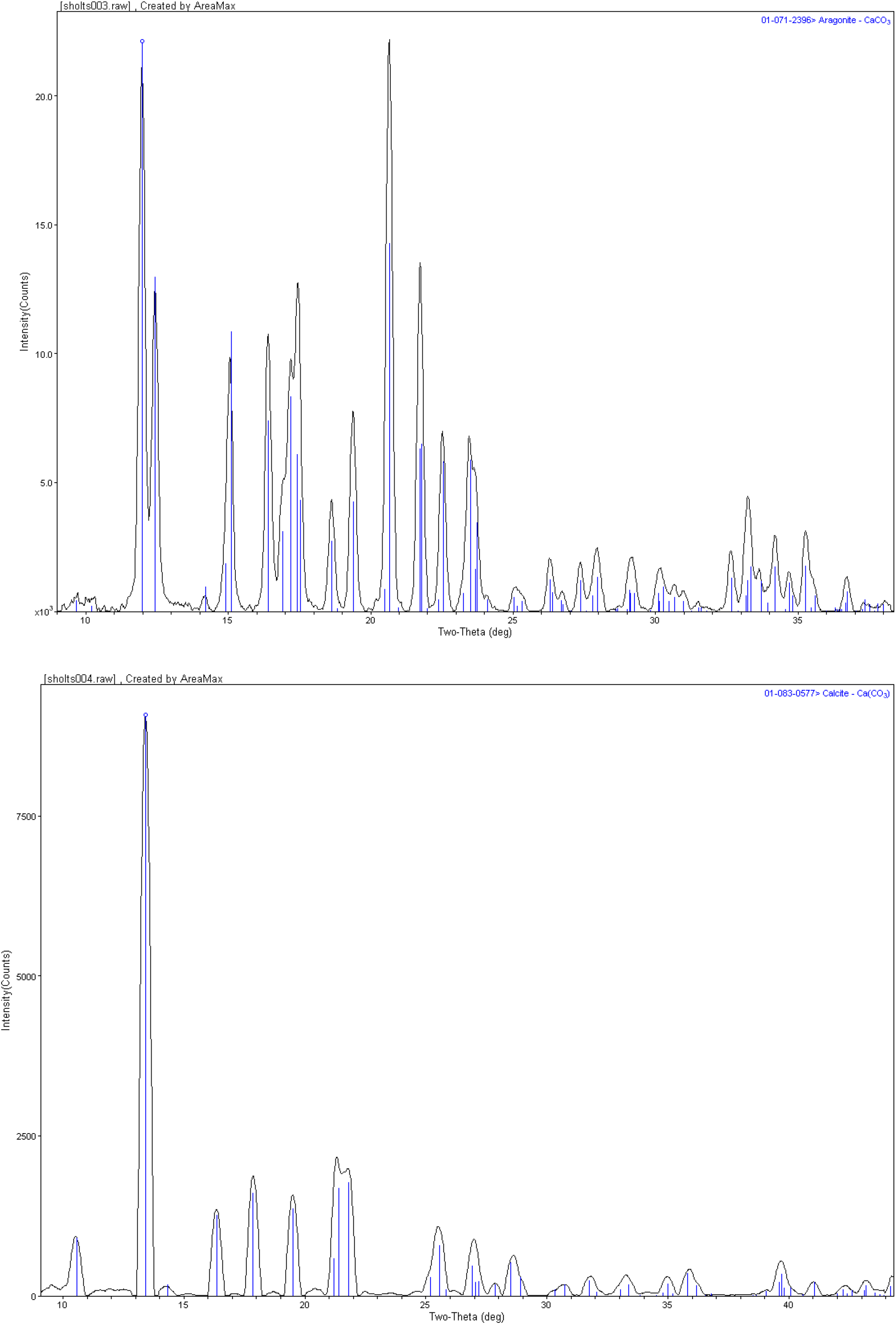
X-ray diffractograms of scrapings from the inner (top) and outer (bottom) layers of modern red abalone shell, perfectly matching reference data for respectively aragonite (ICDD # 01-071-2396) and calcite (ICDD # 01-083-0577).

**Supplementary Figure S7.**
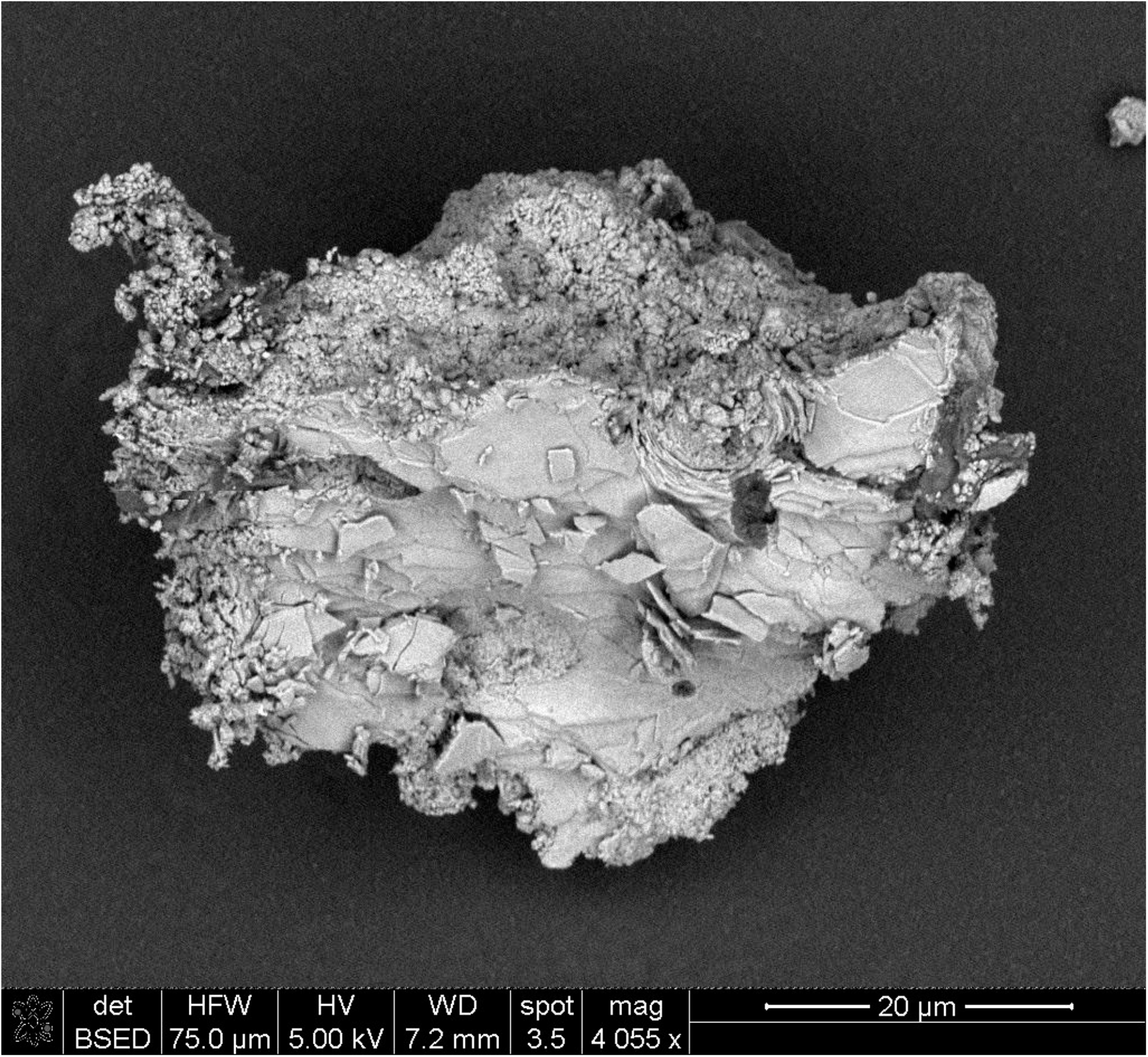
SEM image of a particle from a modern rhizolith collected on San Nicolas Island, showing fragments of aragonite plates in the matrix. Clearly, eroded sea shell material is present in the soil from which the rhizoliths form.

**Supplementary Figure S8.**
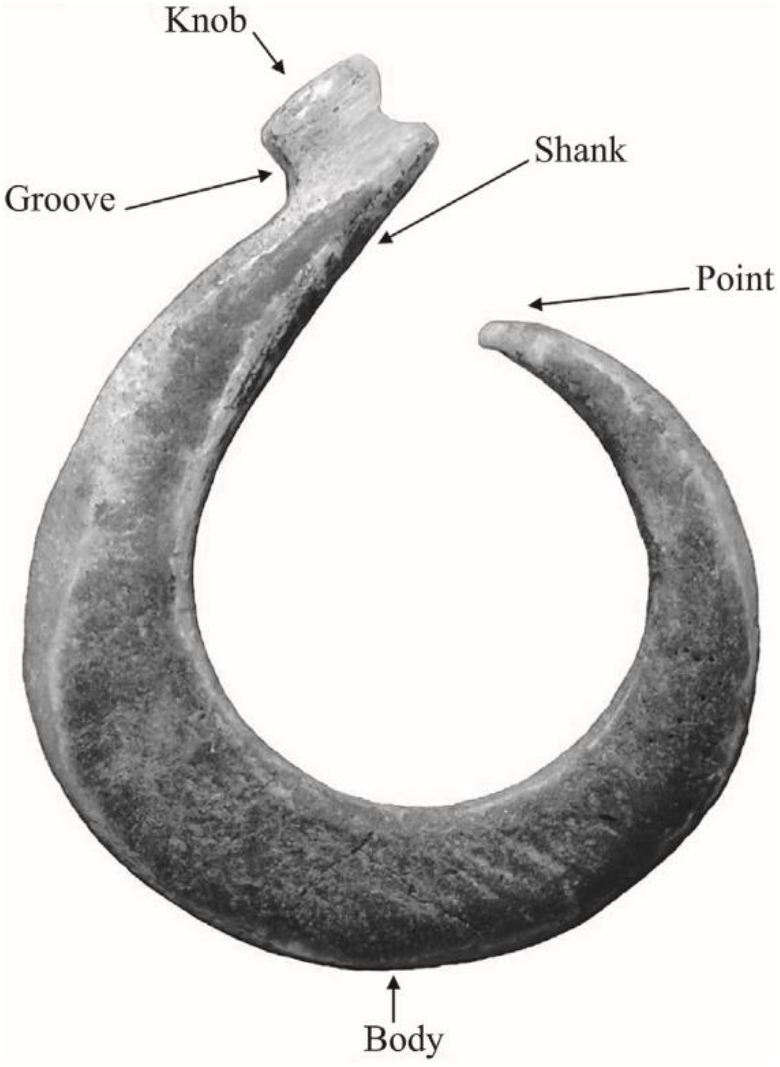
Anatomy of a single piece shell fishhook from the Tule Creek Site. (Adapted from Cannon, A.C., 2006: Giving Voice to Juana María’s People: the Organization of Shell and Exotic Stone Artifact Production and Trade at a Late Holocene Village on San Nicolas Island, California. Unpublished M.A. thesis, Department of Anthropology, Humboldt State University, Arcata.)

**Supplementary Figure S9.**
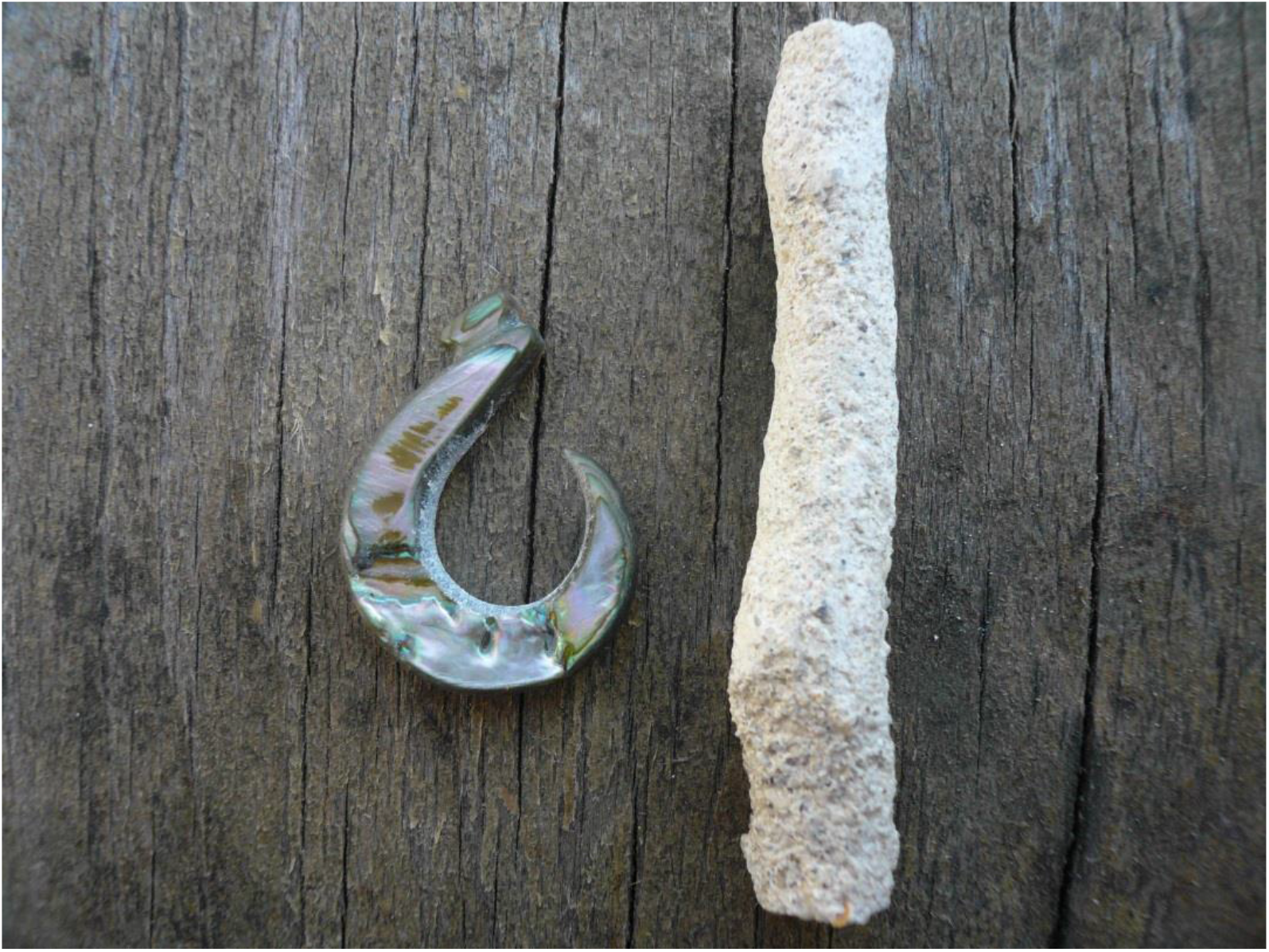
Left: Modern replica of the kind of circular fishhooks found on San Nicolas Island, crafted from abalone shell. Right: The rhizolith that was used as a “rat tail file” to rasp the inner curvature of the hook. Note how the rasping has smoothed the modern rhizolith, giving it an appearance similar to that of the archaeological rhizoliths.

**Supplementary Figure S10.**
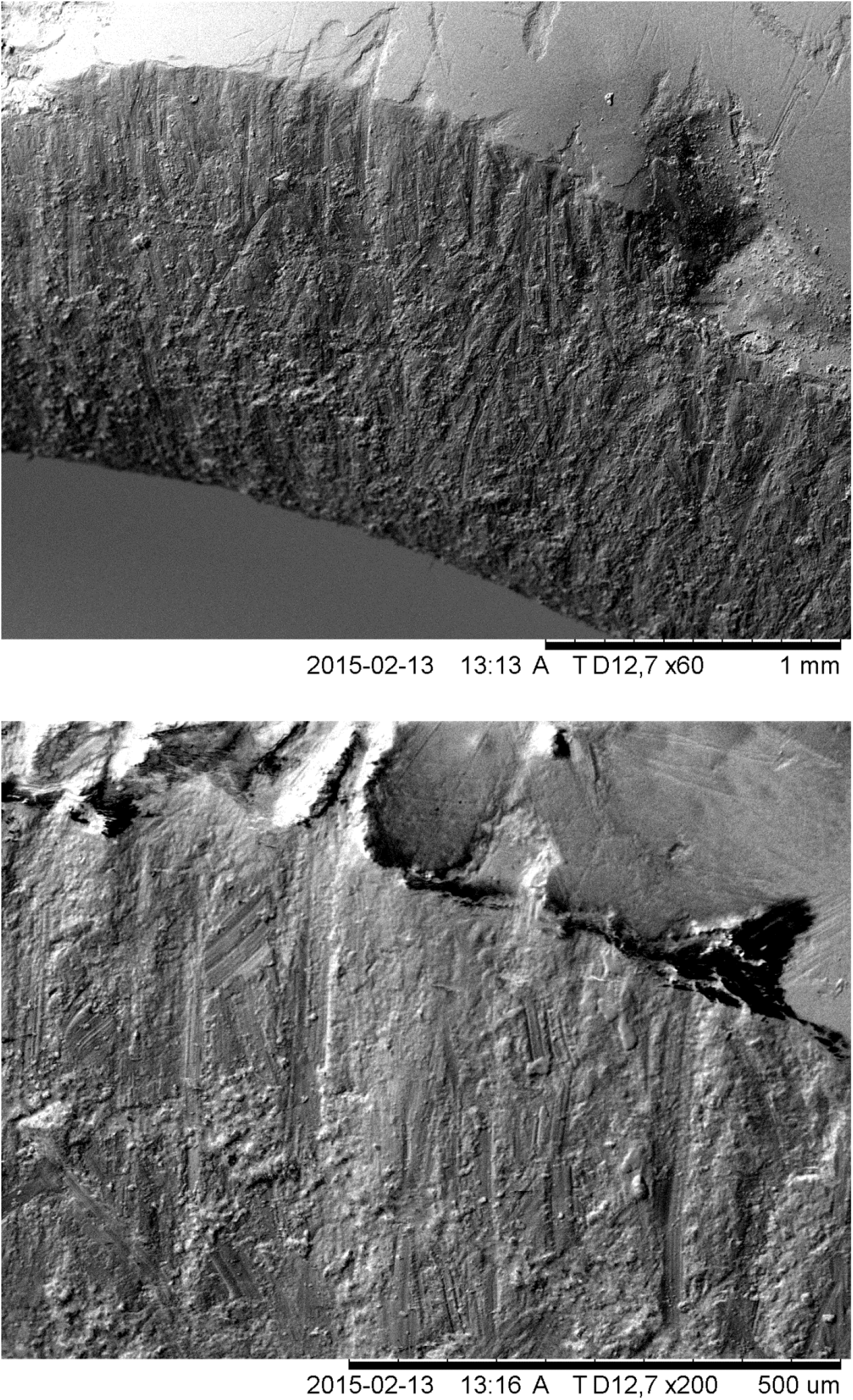
SEM images (top: ×50; bottom ×200) of the inner curvature of a modern fishhook replica, after filing with a rhizolith. Numerous characteristic tool marks are present.

**Supplementary Figure S11.**
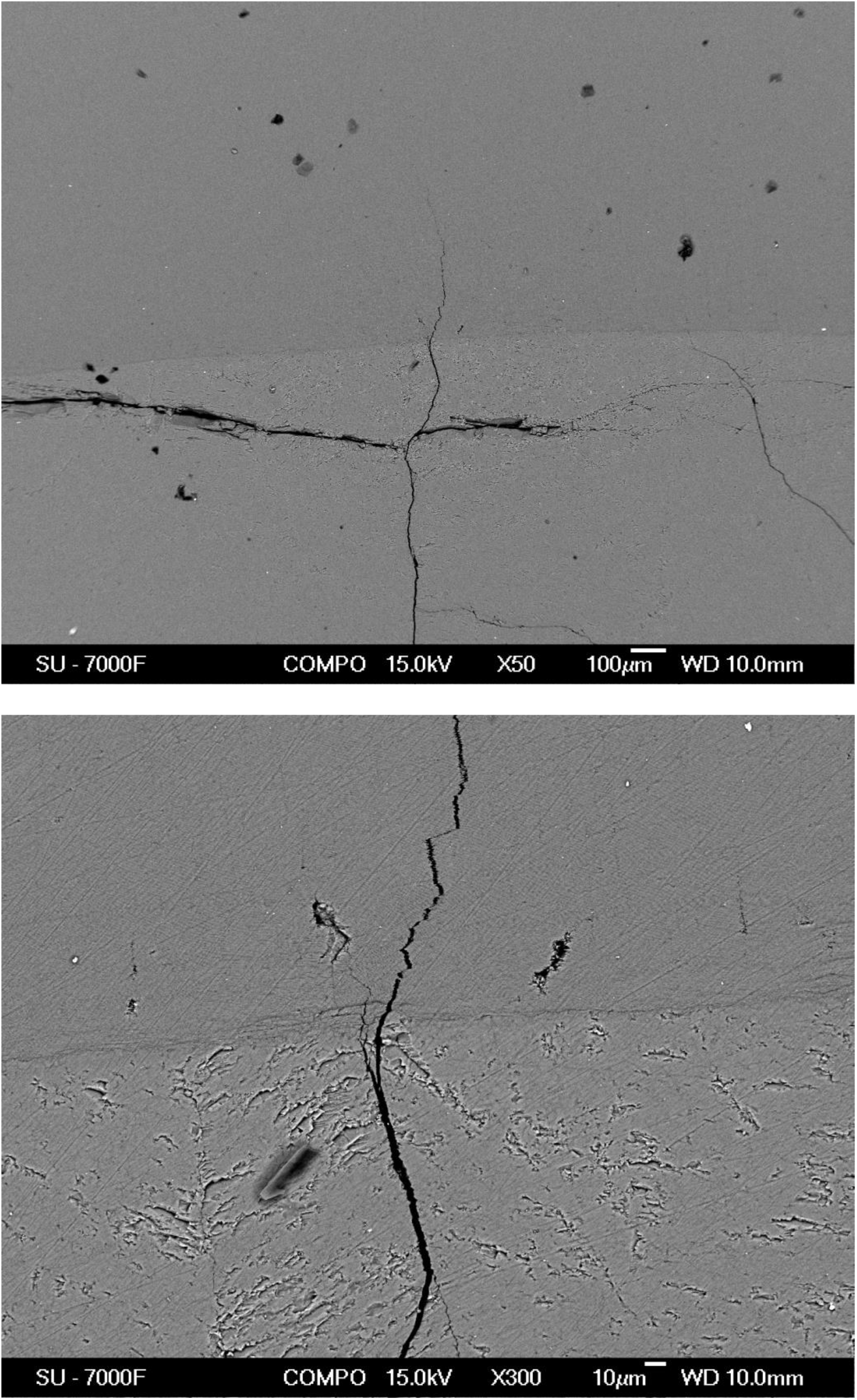
SEM images (top: ×50; bottom ×300) of a cross-section of red abalone shell, showing the intersection between the outer calcitic (lower in image) and inner nacreous (upper in image) parts (compare Fig. 5 in the main article). In the outer calcitic part, multiple cracks run in different directions, but these cracks come to a halt when entering the nacre.

**Supplementary Figure S12.**
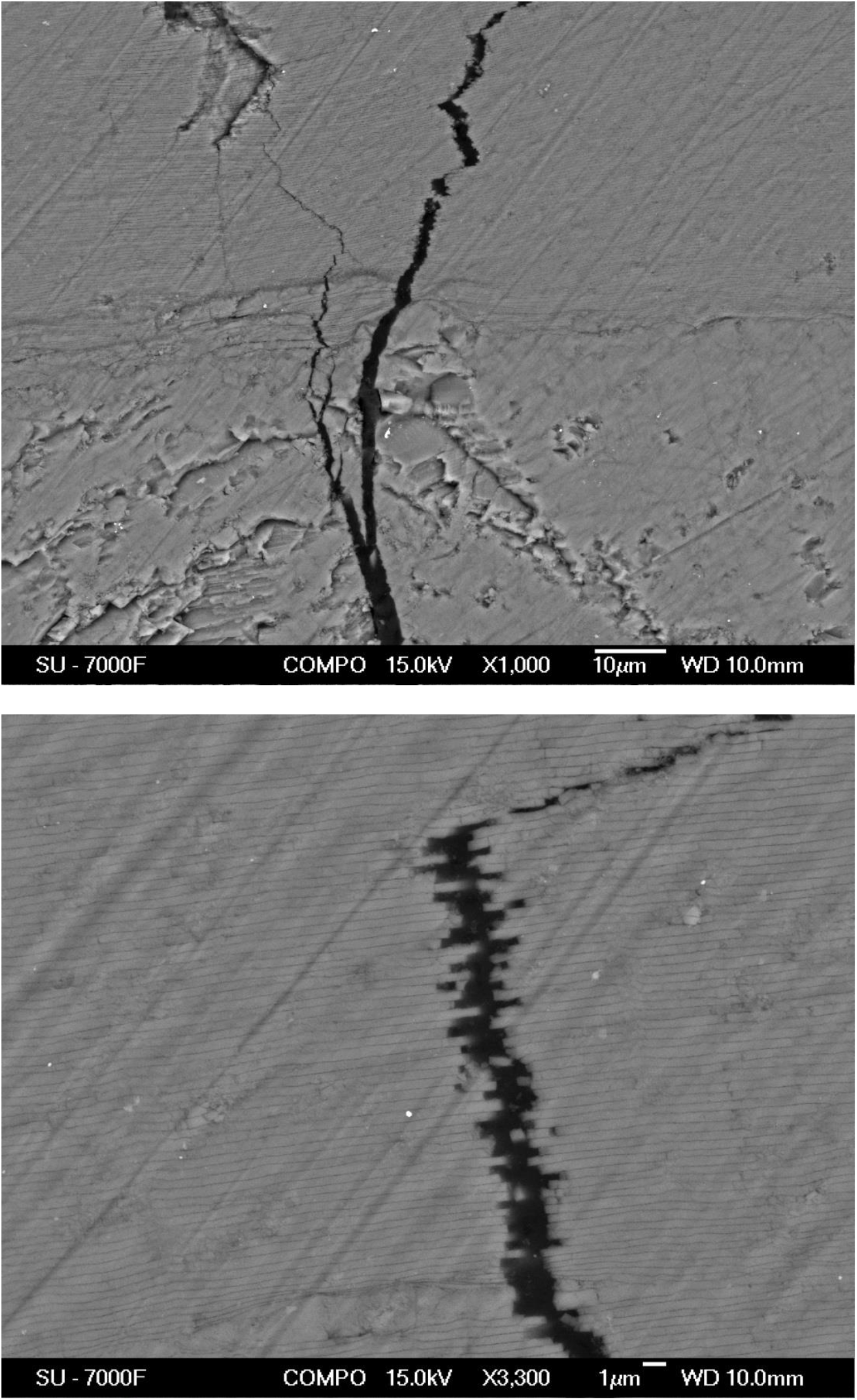
SEM images (top: ×1000; bottom ×3300) of a cross-section of red abalone shell, showing the intersection between the outer calcitic (lower in image) and inner nacreous (upper in image) parts (compare Fig. 5 in the main article). The nacre material consists of layers of aragonite plates in a structural arrangement that impedes crack propagation.

